# Concatenated 16S rRNA Sequence Analysis Improve Bacterial Taxonomy

**DOI:** 10.1101/2022.08.09.503025

**Authors:** Bobby Paul

## Abstract

Microscopic, biochemical, molecular, and computer-based approaches are extensively used to identify and classify bacterial populations. Further, advances in DNA sequencing and bioinformatics workflows facilitated sophisticated genome-based methods for microbial taxonomy. Although sequencing of 16S rRNA gene is widely employed to identify and classify the bacterial community as a cost-effective and single-gene approach. However, the accuracy of the 16S rRNA sequence-based species identification is limited by multiple copies of the gene and their higher sequence identity between closely related species. Availability of a large volume of bacterial whole-genome data provided an opportunity to develop comprehensive species-specific 16S rRNA reference libraries. With defined rules, we have concatenated four 16S rRNA gene copy variants to develop a species-specific reference library. Using this approach, species-specific 16S rRNA gene libraries were developed for four closely related *Streptococcus* species (*S. gordonii, S. mitis, S. oralis*, and *S. pneumoniae*). Sequence similarity and phylogenetic analysis of concatenated 16S rRNA copies yielded better resolution than single gene copy approaches. The approach is very effective to classify genetically related species, and it may reduce misclassification of bacterial species and genome assemblies.

## Introduction

The 16S ribosomal RNA (16S rRNA) encoding region is extensively studied to identify and classify bacterial species. The 16S rRNA is a conserved component of the 30S small subunit of a prokaryotic ribosome. The gene is ∼1500 base pair (bp) long, and it consists of nine variable regions (Reller et al. 2007; Sabat et al. 2017). For decades, the sequence of 16S rRNA has been used as a potential molecular marker in culture-independent methods to identify and classify diverse bacterial communities (Clarridge, 2004; Johnson et al. 2019). The 16S rRNA sequences are currently being used as an accurate and rapid method to study bacterial evolution, phylogenetic relationships, populations in an environment, and quantification of abundant taxa (Vetrovsky and Baldrian, 2013; Srinivasan et al. 2015; Peker et al. 2019).

Despite the wide range of applications, few shortcomings limit the accuracy of results derived through the 16S rRNA sequence analysis. One such aspect is that the 16S rRNA gene has poor discriminatory power at the species level (Winand et al. 2020), and the copy number can vary from 1 to 15 or even more (Vetrovsky and Baldrian, 2013; Winand et al. 2020). The presence of multiple variable copies of this gene makes distinct data for a species. Hence, gene copy normalization (GCN) is necessary prior to the sequence analysis. However, studies show that the GCN approach does not improve the 16S rRNA sequence analyses in real scenarios and suggests a comprehensive species-specific catalogue of gene copies (Starke et al. 2021). Secondly, the intra-genomic variations between the 16S rRNA gene copies were observed in several bacterial genome assemblies (Paul et al. 2019). Only a minority of the bacterial genomes harbor identical 16S rRNA gene copies, and sequence diversity increases with increasing copy numbers (Vetrovsky and Baldrian, 2013). Further, currently available 16S rRNA-based bioinformatics approaches are not always amenable to classify bacterium at the species level due to high inter-species sequence similarities (Peker et al. 2019; Deurenberg et al. 2017).

A few other issues are also related to the sequencing and bioinformatics analysis of 16S rRNA gene regions. These include the purity of bacterial isolates, the quality of isolated DNA, and the possibility of chimeric molecules (Janda and Abbott, 2007; Church et al. 2020). Base-call errors can also mislead the sequence identity and phylogenetic inferences (Alachiotis et al. 2013). The other concerns on sequence-based analysis, comparison, and species identification include the number of base ambiguities processed, gaps generated during sequence comparison, and algorithm (local or global) used for the sequence alignment. The local alignment algorithm is extensively used for sequence similarity based species identification. Several studies were conducted to identify the best variable region or combination of variable regions for bacterial classification, and a consensus remains to be implemented (Janda and Abbott, 2007; Johnson et al. 2019; Winand et al. 2020). Usage of misclassified sequence as a reference and improper bioinformatics workflows mislead the bacterial taxonomy. Further, the growth of bioinformatics and genetic data has placed genome-based microbial classification in researchers with little or no taxonomic experience, which may also mislead the bacterial taxonomy (Baltrus, 2016).

A few bacterial identification systems with high resolution have been developed using the sequence of polymerase chain reaction (PCR) amplified ∼4.5 kb long 16S–23S rRNA regions (Benítez-Páez and Sanz, 2017; Sabat et al. 2017; Kerkhof et al. 2017). However, these approaches have a few limitations, such as the lack of reference 16S–23S rRNA sequence databases and complementary bioinformatics resources for reliable species identification (Sabat et al. 2017). The recent advancements in bioinformatics workflows (Winand et al. 2020; Schloss, 2020) and reference databases such as SILVA, EzBioCloud (Quast et al. 2013; Yoon, 2017) improved 16S rRNA-based bacterial taxonomy. However, a few recent genome-based studies highlighted the misclassification incidences in bacterial species and genome assemblies (Steven et al. 2017; Martínez-Romero, et al. 2018; Mateo-Estrada et al. 2019; Bagheri et al. 2020).

Nowadays, conventional and high throughput sequencers can amplify all the nine variable regions of the 16S rRNA gene. Although, many 16S rRNA-based bacterial identification studies lack a complete set of variable regions (Stackebrandt et al. 2021). The classical and high throughput sequencing technologies produce a large volume of whole-genome data. There is an urgent need to translate the genomic data for convenient microbiome analyses that ensure clinical practitioners can readily understand and quickly implement it (Church et al. 2020). Hence, we intended to demonstrate the workflow to develop species-specific concatenated 16S rRNA reference libraries and its applications. The species-specific libraries can yield better resolution in sequence similarity and phylogeny based bacterial classification approaches.

## Materials and Methods

### Estimation of variations in intra-genomic 16S rRNA gene copies

Sequence alignment of 16S rRNA copies at the intra-genomic level shows a higher degree of variability in species belonging to the *Firmicutes* and *Proteobacteria* (Vetrovsky and Baldrian, 2013; Ibal et al. 2019). Hence, we used eight 16S rRNA copies (Supplementary data 1) retrieved from the whole-genome of *Enterobacter asburiae* strain ATCC 35953 (NZ_CP011863.1). The BLAST (Altschul et al. 1990) and Clustal Omega (Sievers et al. 2011) sequence alignment algorithms were used to estimate intra-genomic variability between the 16S rRNA gene copies. Phylogenetic relatedness between intra-genomic 16S rRNA copies were estimated using the Maximum Likelihood method (Tamura-Nei model; 500 bootstrap replicates) with MEGA X software (Kumar et al. 2018).

### Construction of species-specific concatenated 16S rRNA reference libraries

Previous studies have reported that several bacterial species shares more than 99% sequence identity in the16S rRNA encoding region. Hence, the 16S rRNA-based bacterial identification methods failed to discriminate such genetically related species (Deurenberg et al. 2017; Devanga-Ragupathi et al. 2018). It has been reported that *Streptococcus mitis* and *Streptococcus pneumoniae* are almost indistinguishable from each other based on the sequence similarity of their 16S rRNA regions (Reller et al. 2007; Lal et al. 2011). To develop species-specific barcode reference libraries, the study used 16S rRNA gene copies from whole-genome assemblies of four closely related species of *Streptococcus* (*S. gordonii, S. mitis, S. oralis* and *S. pneumoniae*).

More than 385000 whole-genome assemblies are currently available for prokaryotes at the Genome database (https://www.ncbi.nlm.nih.gov/genome). Most microbial genomes were sequenced with high throughput sequencing technologies such as Illumina/Ion-Torrent (short read sequencing) and PacBio/Nanopre (long read sequencing). Further, many of these whole-genome assemblies are derived through a hybrid assembly of short and long read sequence data. The large volume of high throughput data can be effectively used to develop advanced genome based approaches for microbial systematics. The genomic data is available in four assembly completion levels (contig, scaffold, chromosome, and complete). We used only the genomes assemblies in the ‘complete’ stage to retrieve 16S rRNA gene copies.

The study retrieved full-length 16S rRNA gene copies from 16 genome assemblies belonging to four *Streptococcus* species (*S. gordonii, S. mitis, S. oralis*, and *S. pneumoniae*). The detailed information on the dataset used to develop species-specific concatenated reference libraries is provided in Table 2 and the sequences are provided in Supplementary data 2. To maintain the equal length, sequences were trimmed out beyond the universal primer pair fD1 - 5’-GAG TTT GAT CCT GGC TCA-3’ and rP2 - 5’-ACG GCT AAC TTG TTA CGA CT-3’ (Weisburg et al. 1991) for full-length *in silico* 16S rDNA amplification. We used MEGA X software to perform multiple sequence alignment and identify the intra-species Parsimony informative (Parsim-info) variable sites. A species-specific barcode reference library covering entire Parsim-info variable sites was constructed by concatenating four 16S rRNA gene copies representing four different strains of a species. The rationale behind the selection of four copies for a species-specific barcode reference library is: (i) a maximum of four variations can be found on a single site, and (ii) earlier studies have shown that the mean 16S rRNA copies per genome is four (Vetrovsky and Baldrian, 2013).

### Demonstration of concatenated 16S rRNA in sequence similarity and phylogeny

We discussed a few case studies to demonstrate the classical sequence similarity and phylogeny-based approaches using concatenated species-specific 16S rRNA reference sequence libraries. The study selected nine Sanger sequenced 16S rRNA gene shown higher sequence similarity with multiple species of *Streptococcus*. Web-based BLAST2 program for aligning two or more sequences was used to estimate the maximum score, total alignment score, and sequence identity. Single copy of the 16S rRNA region derived through Sanger sequencing or retrieved from a whole-genome assembly can be considered as ‘Query sequence’. The concatenated species-specific reference libraries must be provided in the ‘Subject sequence’ section. To perform an accurate phylogenetic analysis, it is mandatory that the target sequence (length=n bp) have to be concatenated four times (length=n×4 bp), appending next to the last base. Phylogenetic relatedness was estimated using the Maximum Likelihood method (Tamura-Nei model; 500 bootstrap replicates) with MEGA X software.

## Results

### Intra-genomic 16S rRNA variations in *Enterobacter asburiae*

Historically, sequences of the 16S rRNA gene were used to identify known and new bacterial species. However, this method is impacted by several factors such as amplification efficiency, poor discriminatory power at the species level, multiple polymorphic 16S rRNA gene copies, and improper bioinformatics workflows for the data analysis. The genome have eight 16S rRNA gene copies that showed a mean identity of 99.29% in sequence alignment using Clustal Omega (global alignment), whereas BLAST (local alignment) analysis resulted in an average of 99% identity between the copies (Table 1). Hence, the selection of an appropriate algorithm have a significant role in the estimation of percent identity, and a vital role in sequence-based species delineation. Global sequence alignment programs generally perform better for highly identical sequence pairs, and the algorithm considers all the bases for the estimation of sequence identity. The multiple sequence alignment showed 22 variable sites in 16S rRNA gene copies of *E. asburiae* genome (Fig. 1).

**Table 1.** Percent identity of eight intra genomic 16S rRNA regions from *Enterobacter asburiae* strain ATCC 35953 (NZ_CP011863.1). Percent identity given below the diagonal line is calculated with Clustal Omega software (Mean identity: 99.29%) and those above the diagonal line was calculated with BLASTN program (Mean identity: 99.00%). Genome coordinates of 16S rRNA copies-R1: 2686082-2687660 (1579 bp); R2: 3148265-3149814 (1550 bp); R3: 3313470-3315019 (1550 bp); R4: 3583942-3585481 (1540 bp); R5:3684745-3686294 (1550 bp); R6: 3771751-3773300 (1550 bp); R7: 3968538-3970087 (1550 bp); R8: 4647650-4649199 (1550 bp)

| 16S rRNA copies | R1 | R2 | R3 | R4 | R5 | R6 | R7 | R8 |
| --- | --- | --- | --- | --- | --- | --- | --- | --- |
| **R1** |  | 98.10 | 98.04 | 97.47 | 98.04 | 97.47 | 97.59 | 98.04 |
| **R2** | 99.10 |  | 99.74 | 99.23 | 99.94 | 99.29 | 99.48 | 99.94 |
| **R3** | 98.97 | 99.74 |  | 99.23 | 99.68 | 99.03 | 99.23 | 99.81 |
| **R4** | 98.90 | 99.41 | 99.41 |  | 99.16 | 98.52 | 98.71 | 99.29 |
| **R5** | 99.03 | 99.94 | 99.68 | 99.35 |  | 99.23 | 99.42 | 99.87 |
| **R6** | 98.39 | 99.29 | 99.03 | 98.70 | 99.23 |  | 99.68 | 99.23 |
| **R7** | 98.58 | 99.48 | 99.23 | 98.89 | 99.42 | 99.68 |  | 99.42 |
| **R8** | 99.03 | 99.94 | 99.81 | 99.48 | 99.87 | 99.23 | 99.42 |  |

**Fig. 1.**
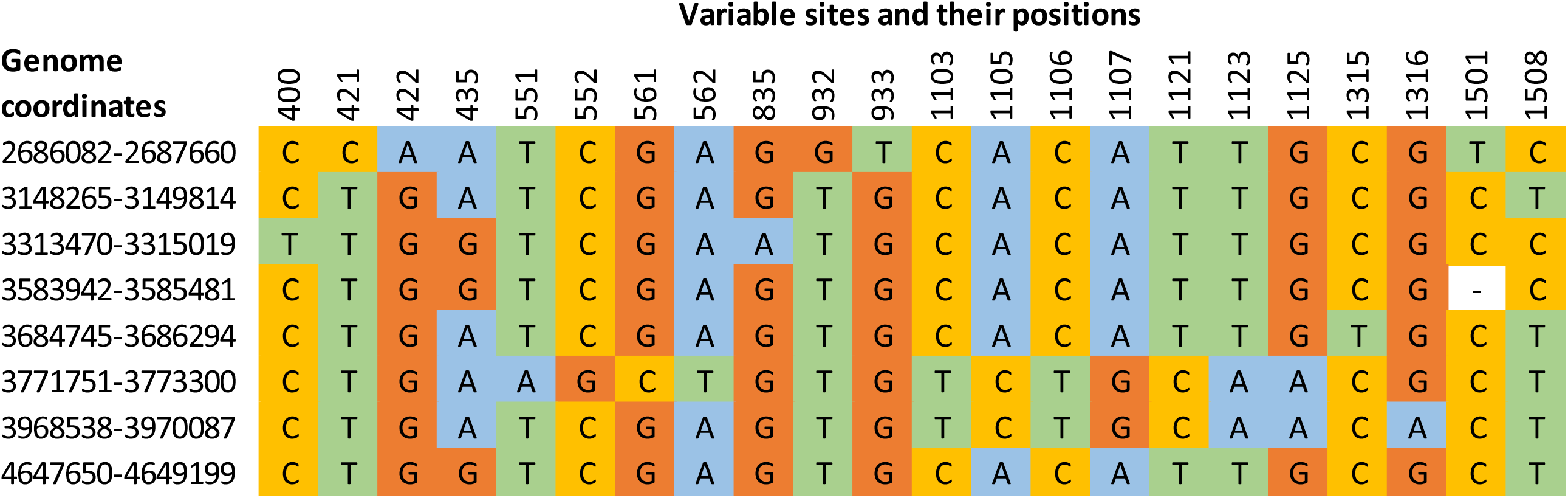
Multiple sequence alignment of eight intra genomic 16S rRNA gene copies from *Enterobacter asburiae* strain ATCC 35953 (NZ_CP011863.1) showing 22 variable sites.

The evolutionary relationship between species is usually represented in a phylogenetic tree drawn using a single barcode gene, multiple genes, or whole genomes. However, bacterial species nomenclature is mainly designated based on the confidence obtained from the phylogenetic tree derived through single copy 16S rRNA analysis. To highlight how the intra-genomic variations of 16S rRNA copies influence the single gene phylogeny for species delineation. We constructed a phylogenetic tree using eight 16S rRNA gene copies of *E. asburiae* reference genome showing multiple nodes (Fig. 2). The sequence similarity and phylogeny-based analysis indicate that the intra-genomic variations in 16S rRNA copies may mislead the bacterial taxonomy in single gene copy approaches.

**Fig. 2.**
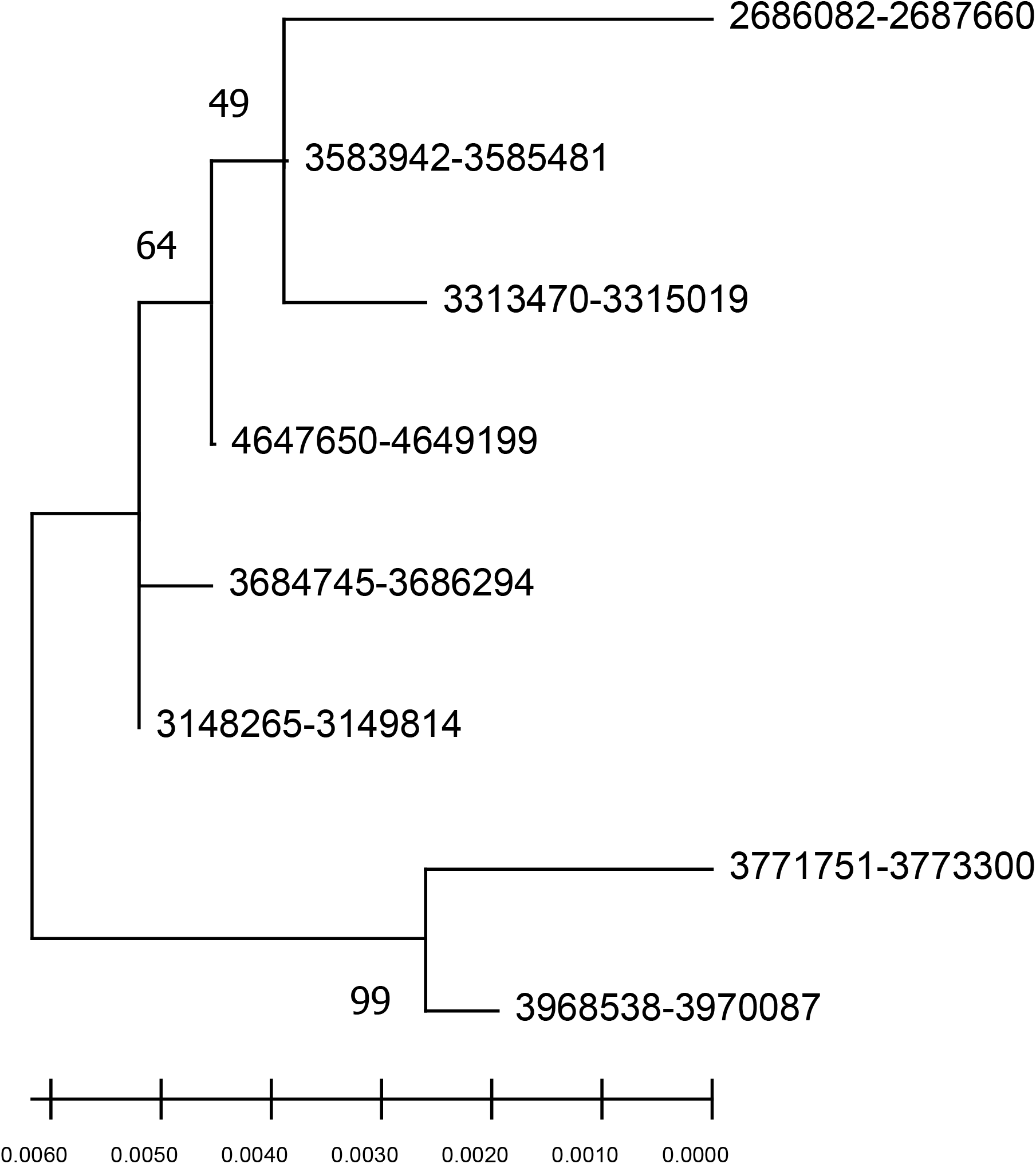
Phylogenetic tree of eight intra genomic 16S rRNA gene copies from *Enterobacter asburiae* strain ATCC 35953 (NZ_CP011863.1). The node label denotes the coordinate of 16S rRNA regions in the genome.

### Species-specific concatenated 16S rRNA libraries

We selected four *Streptococcus* species (*S. gordonii, S. mitis, S. oralis*, and *S. pneumoniae*) to construct species-specific concatenated 16S rRNA reference libraries. The study used four whole genome assemblies in the ‘complete’ stage to construct a species-specific barcode library. Four copies of 16S rRNA gene is required to construct the concatenated library for a species. The details of constructed species-specific libraries is listed in Table 2 and the sequence is provided in Supplementary data 3. The 16S rRNA sequence analysis shows 24 Parsim-info variable sites for *S. oralis*, 11 variations in *S. mitis*, seven variations in *S. gordonii*, and six variations found in *S. pneumoniae*.

**Table 2.** Details of whole genome assemblies used for the development of concatenated 16S rRNA reference libraries. One copy of 16S rRNA gene from each strain is used for the concatenation.

| Species | Strains | Genome accession number | No. of 16S rRNA gene copies | Sequencing platform | Species-specific library name | Library length (bp) | No. of Parsim-info sites |
| --- | --- | --- | --- | --- | --- | --- | --- |
| *S. gordonii* | FDAARGOS 1454 | CP077224.1 | 4 | PacBio; Illumina | *S.gordonii*-Ref-I | 6076 | 7 |
|  | NCTC7868 | LR134291.1 | 4 | PacBio |  |  |  |
|  | KCOM 1506 | CP012648.1 | 5 | Illumina |  |  |  |
|  | NCTC9124 | LR594041.1 | 4 | PacBio |  |  |  |
| *S. mitis* | B6 | NC_013853.1 | 4 | NA | *S.mitis*-Ref-I | 6033 | 11 |
|  | KCOM 1350 | CP012646.1 | 3 | Illumina |  |  |  |
|  | SVGS 061 | CP014326.1 | 4 | PacBio; Illumina |  |  |  |
|  | NCTC 12261 | CP028414.1 | 4 | PacBio |  |  |  |
| *S. oralis* | NCTC 11427 | LR134336.1 | 4 | PacBio | *S.oralis*-Ref-I | 6038 | 24 |
|  | 34 | CP079724.1 | 4 | Illumina; Nanopore |  |  |  |
|  | FDAARGOS 886 | CP065706.1 | 4 | PacBio; Illumina |  |  |  |
|  | F0392 | CP034442.1 | 4 | PacBio |  |  |  |
| *S. pneumoniae* | 475 | CP046355.1 | 4 | PacBio | *S.pneumoniae*-Ref-I | 6032 | 6 |
|  | NU83127 | AP018936.1 | 4 | Nanopore; Illumina |  |  |  |
|  | NCTC7465 | LN831051.1 | 4 | PacBio |  |  |  |
|  | 6A-10 | CP053210.1 | 4 | PacBio |  |  |  |

The study used full-length 16S rRNA copies from four different strains to highlight the variations at the species level. The observed intra-species Parsim-info variable sites reside on both conserved and variable regions of 16S rRNA gene. Species-specific concatenated 16S rRNA reference library can be developed with limited number of variable regions. Intra-species variation on 16S rRNA gene copies influences the sequence based bacterial taxonomy. Hence, concatenated 16S rRNA approach yield better resolution than single copy analysis in classical sequence similarity and phylogeny based species identification approaches.

### Demonstration of concatenated 16S rRNA based species identification

The study compared nine 16S rRNA sequences representing *Streptococcus* species (Table 3) with species-specific concatenated reference libraries. Concatenated sequence analysis gives better resolution in sequence similarity search and phylogenetics analysis. The sequence accession numbers GU470907.1 and KF933785.1 classified as *S. mitis* showed a higher maximum and total alignment score with *S. oralis* than *S. mitis* (Table 3). Whereas the sequence (OM368574.1; classified as *S. mitis*) showed a higher sequence alignment score with *S. pneumoniae*. The Fig. 3A shows a Maximum Likelihood tree of the nine 16S rRNA gene sequences with four concatenated species-specific reference libraries. The concatenated GU470907.1 and KF933785.1 sequences showed phylogenetic relationship with *S. oralis* and sequence OM368574.1 genetically related with *S. pneumoniae*. These results indicate that the species-specific concatenated 16S rRNA reference libraries have great potential in the taxonomy of genetically related species. Hence, the study suggests the usage of concatenated variable 16S rRNA copies for sequence similarity and phylogeny-based species identification. Species-specific reference library with concatenated 16S rRNA gene copies provides better resolution in phylogenetic analysis than the single copy inference.

**Table 3.** Similarity of selected sequences against the concatenated species-specific 16S rRNA reference libraries.

| GenBank Acc. No. | Species | *S. gordonii*-Ref-I |  |  | *S. mitis*-Ref-I |  |  | *S. oralis*-Ref-I |  |  | *S. pneumoniae*-Ref-I |  |  |
| --- | --- | --- | --- | --- | --- | --- | --- | --- | --- | --- | --- | --- | --- |
|  |  | Max Score | Total Score | Identity (%) | Max Score | Total Score | Identity (%) | Max Score | Total Score | Identity (%) | Max Score | Total Score | Identity (%) |
| AJ295848.1 | *S. mitis* | 2495 | 9967 | 96.45 | 2769 | 11027 | 99.80 | 2758 | 10851 | 99.67 | 2752 | 10982 | 99.60 |
| AM157428.1 | *S. mitis* | 2462 | 9845 | 96.05 | 2724 | 10866 | 99.27 | 2702 | 10685 | 99.01 | 2708 | 10805 | 99.07 |
| NR_028664.1 | *S. mitis* | 2499 | 9991 | 96.45 | 2776 | 10979 | 99.87 | 2750 | 10864 | 99.54 | 2724 | 10888 | 99.27 |
| GU470907.1 | *S. mitis* | 2536 | 10096 | 96.91 | 2715 | 10796 | 99.14 | 2787 | 10936 | 100 | 2091 | 10716 | 98.87 |
| KF933785.1 | *S. mitis* | 2466 | 9832 | 96.06 | 2667 | 10593 | 98.54 | 2673 | 10650 | 98.61 | 2632 | 10502 | 98.15 |
| OM368574.1 | *S. mitis* | 2475 | 9896 | 96.24 | 2754 | 10968 | 99.67 | 2732 | 10814 | 99.40 | 2760 | 10990 | 99.73 |
| OM368578.1 | *S. pneumoniae* | 2475 | 9896 | 96.24 | 2754 | 10968 | 99.67 | 2732 | 10814 | 99.40 | 2760 | 10990 | 99.73 |
| AM157442.1 | *S. pneumoniae* | 2470 | 9863 | 96.12 | 2702 | 10779 | 99.01 | 2715 | 10726 | 99.14 | 2702 | 10777 | 99.01 |
| NR_117719.1 | *S. oralis* | 2531 | 10074 | 96.84 | 2710 | 10774 | 99.07 | 2787 | 10925 | 100 | 2697 | 10739 | 98.94 |

**Fig. 3.**
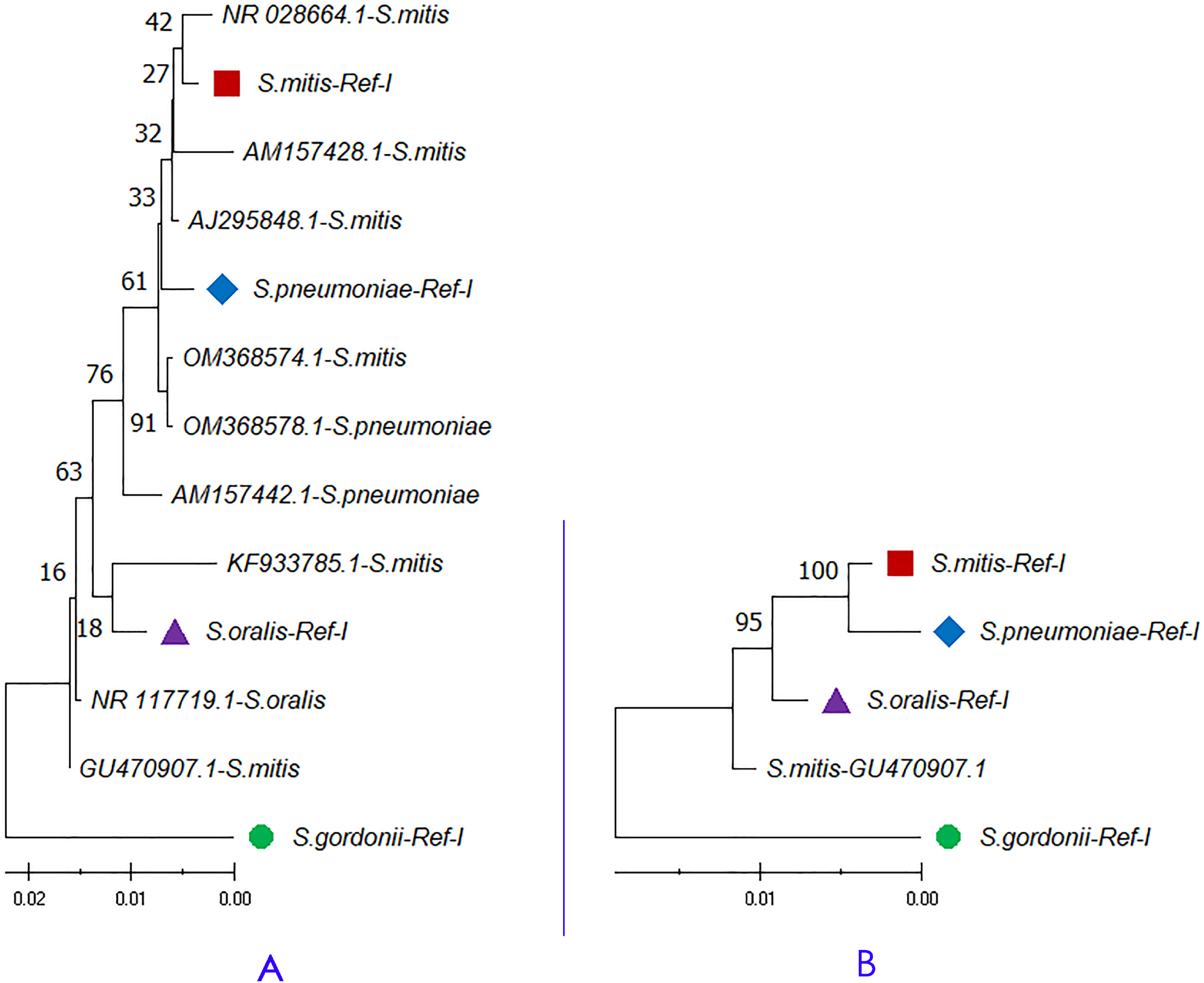
A) Phylogenetic analysis of analysis of randomly selected 16S rRNA sequences classified as *Streptococcus* species. B) Concatenated 16S rRNA phylogeny of *Streptococcus mitis* sequence (Ac. No. GU470907.1) showed 100% identity with *Streptococcus oralis* genome (Ac. No. CP034442.1) in BLAST based sequence similarity search. The node name highlighted in shapes (●, ■, ▲, ◆) represents the four species-specific reference libraries.

## Discussion

The 16S rRNA encoding region sequences are considered a conventional and robust method for identifying the bacterial species. The barcode gene is widely used in sequence similarity, phylogeny, and metagenome based species identification. Recently, Church et al. (2020) reviewed in detail the Sanger sequencing of 16S rRNA gene, sequence data analysis, and result interpretation. However, discrimination of closely related species identification through sequences of 16S rRNA gene is a challenge, and it may lead to species misidentifications (Boudewijns et al. 2006). The 16S rRNA gene copies can vary from 1 to 15 in a genome, and the copy number of variations is taxon-specific (Vetrovsky and Baldrian, 2013). The 16S rRNA sequence variation found at intra-genomic level and between the strains of a species as well. Sequence diversity increases with the increasing 16S rRNA copy numbers. About 15% of the genomes have only a single copy of the 16S rRNA gene, and only a minority of bacterial genomes harbours identical 16S rRNA gene copies (Vetrovsky and Baldrian, 2013). Amplification of limited number of variable regions cannot achieve the taxonomic resolution achieved by sequencing the entire gene (Johnson et al. 2019). Usage of misclassified 16S rRNA sequences as a reference and inappropriate bioinformatics workflows also mislead the taxonomic assignment.

Several bioinformatics resources are extensively used for the 16S rRNA sequence analysis and bacterial identification. However, several researchers report the sequence similarity derived through a local alignment algorithm. Earlier reports have suggested that the species belonging to the taxa Gammaproteobacteria show higher intra-species variability (Vetrovsky and Baldrian, 2013). Hence, we estimated the percent identity of intra-genomic 16S rRNA gene copies of *Enterobacter asburiae* using local and global alignment algorithms. The reference genome of *E. asburiae* has eight 16S rRNA gene copies in its genome. The BLAST and Clustal sequence alignment algorithms yielded marginally varying results for the intra-genomic 16S rRNA gene copies. Local alignment algorithms may not consider base mismatches at the sequence ends for calculating percent identity, while global alignment algorithms consider entire bases. Therefore, we suggest that global sequence alignment is best for estimating intra and inter-species identity for single gene copies. However, BLAST can calculate the total alignment score with multiple paralogues regions. Hence, we suggest BLAST2 for estimating the sequence similarity using concatenated barcode reference libraries.

The GenBank (Leray et al. 2019) and NCBI 16S database for bacteria (Winand et al. 2020) are reliable for species-level identification and classification. However, few earlier studies have been highlighted the misclassification of species and genome assemblies at public genetic databases (Parks et al. 2018; Varghese et al. 2015). For example, the 16S rRNA sequence (Ac. No. LT707617) shows the organism as *Streptococcus mitis*. Conventional BLAST-based sequence similarity search shows the highest identity of 99.60% with *S. mitis* 16S rRNA sequence (Ac. No. AB002520). However, the 16S rRNA sequence (Ac. No. LT707617) did not show significant similarity with other 16S rRNA reference sequences available for *S. mitis*. Further, the sequence also shows 99.44% identity with reference 16S rRNA sequences of *S. gordonii*. Hence, we performed a sequence alignment of the sequence (Acc. No. LT707617) against species-specific concatenated 16S rRNA reference libraries for *S. gordonii* (*S*.*gordonii-*Ref-I), and *S. mitis* (*S*.*mitis-*Ref-I). The alignment resulted in a significant identity of 99.44% with *S*.*gordonii-*Ref-I (2279 maximum and 9041 total alignment score) than *S*.*mitis*-Ref-I (97.13% identity with 2119 maximum and 8449 total alignment score). Single copy BLAST results may show only a minor fraction of the difference in percent identity and maximum or total alignment score for closely related species. However, sequence similarity estimation using species-specific concatenated reference libraries shows a significant difference in total alignment score, as it is aligned against four copies. Hence, 16S rRNA analysis with species-specific concatenated barcode reference library will give better accuracy for bacterial classification than approaches using a single copy.

Several 16S rRNA sequences show 100% identity with multiple species, which is the major challenge in sequence-based species identification. For example, 16S rRNA sequence from *S. mitis* sequence (Ac. No. GU470907.1; 1522 bp) share 100% identity with 16S rRNA gene from *S. oralis* strain ATCC 35037 genome (Ac. No. CP034442.1). Hence, we compared the sequence (GU470907.1) against the species-specific concatenated reference libraries for *S. oralis* (*S*.*oralis-*Ref-I), and *S. mitis* (*S*.*mitis-*Ref-I). The result showed 100% identity with *S. oralis* (2787 maximum and 10936 total alignment score), and 99.14% identity with *S. mitis* (2715 maximum and 10796 total alignment score). Further, we plotted a phylogenetic tree of GU470907 (1509 × 4 = 6036 bp) with reference libraries *S*.*mitis-*Ref-I, and *S*.*oralis-* Ref-I. The Maximum Likelihood-based phylogenetic tree showed that the *S. mitis* (GU470907.1) sequence is closely related to *S. oralis* than *S. mitis* (Fig. 3B). Concatenated 16S rRNA-based estimation of sequence similarity and a phylogenetic inference provides resolution than single-gene approaches. These results show that concatenated 16S rRNA approach is very effective in discriminating genetically related bacterial species. Further, other studies also highlighted that the phylogenetic tree inferred from vertically inherited protein sequence concatenation provided higher resolution than those obtained from a single copy (Ciccarelli et al. 2006; Thiergart et al. 2014).

Recent phylogenetic studies using concatenated multi-gene sequences data highlighted the importance of incorporating variation in gene histories and which will improve the traditional phylogenetic inferences (Devulder et al. 2005; Johnston et al. 2019). As a cost-effective approach, we combine substantial variations in 16S rRNA gene copies from a species to examine the performance of the single gene concatenation approach. Analysis using concatenated 16S rRNA gene approach have some advantages: (i) the gene is present in all the bacterial species, (ii) the gene is weakly affected by horizontal gene transfer, (iii) the approach is very cost-effective, (iv) large volume of reference genomic data available for several bacterial species, (v) effective to discriminate closely related bacterial species, and (vi) availability of bioinformatics resources for data analytics.

Sequencing and analysis of the 16S rRNA gene region is considered the gold standard for identifying and classifying the bacterial population. The accuracy of bacterial taxonomy based on 16S rRNA barcode regions is limited by the intra-genomic heterogeneity of multiple 16S rRNA gene copies and significant sequence identity of this gene between the closely related taxa. Overcoming these challenges, clinical laboratories are looking forward to translating high throughput microbial genomic data into meaningful, actionable information that clinicians can readily understand and quickly implement for bacterial identification. We suggest not to rely upon one type of analysis, instead and to a certain extent, integrated bioinformatics approaches can avoid misclassification. Developing a species-specific catalogue of concatenated 16S rRNA gene copies for the sequence similarity and phylogenetic studies will give better inference and can be used even in mapping-based metagenome approaches.

## Conclusion

The concatenated 16S rRNA analysis drew the following suggestions:

- Full-length 16S rRNA gene amplification provides better accuracy than inference from a partial gene with a limited number of variable sites.
- Prior to the analysis, trim the bases beyond the primer ends and correct the base-call errors, which will avoid several mismatches in the sequence alignment.
- Estimation of mean 16S rRNA identity at the intra-species level helps to classify the species having a higher degree of intra-genomic 16S rRNA heterogeneity.
- Use full-length 16S rRNA gene copies from whole-genome assemblies (in ‘complete’ stage) rather than partial sequences available at the public genetic databases to construct species-specific concatenated 16S rRNA libraries and further downstream analysis.
- Distinct four 16S rRNA gene copies cover all the Parsim-Info variable sites of a species can be used to construct concatenated species-specific reference library.
- The total alignment score can be considered, if the query sequence shows more or less the same percent identity with multiple species.
- Do not rely only on sequence similarity; make a final decision based on the phylogenetic inference.

## References

Alachiotis N, Vogiatzi E, Pavlidis P, Stamatakis A (2013) Chromatogate: a tool for detecting base mis-calls in multiple sequence alignments by semi-automatic chromatogram inspection. Comput Struct Biotechnol J 6: e201303001. https://doi.org/10.5936/csbj.201303001.

Altschul SF, Gish W, Miller W, Myers EW, Lipman DJ (1990) Basic local alignment search tool. J Mol Biol 215: 403–410. https://doi.org/10.1016/S0022-2836(05)80360-2.

Bagheri H, Severin AJ, Rajan H (2020) Detecting and correcting misclassified sequences in the large-scale public databases. Bioinformatics 36: 4699–4705. https://doi.org/10.1093/bioinformatics/btaa586.

Baltrus DA (2016) Divorcing strain classification from species names. Trends Microbiol 24: 431–439. https://doi:10.1016/j.tim.2016.02.004.

Benitez-Paez A, Sanz Y (2017) Multi-locus and long amplicon sequencing approach to study microbial diversity at species level using the MinIONTM portable Nanopore sequencer. Gigascience 6: 1–12. https://doi.org/10.1093/gigascience/gix043.

Boudewijns M, Bakkers JM, Sturm PDJ, Melchers WJG (2006) 16S rRNA gene sequencing and the routine clinical microbiology laboratory: A perfect marriage? J Clin Microbiol 44: 3469–3470. https://doi.org/10.1128/JCM.01017-06.

Church DL, Cerutti L, Gürtler A, Griener T, Zelazny A, Emler S (2020) Performance and application of 16S rRNA gene cycle sequencing for routine identification of bacteria in the clinical microbiology laboratory. Clin Microbiol Rev 33: e00053–19. https://doi.org/10.1128/CMR.00053-19.

Ciccarelli FD, Doerks T, von Mering C, Creevey CJ, Snel B, Bork P (2006) Toward automatic reconstruction of a highly resolved tree of life. Science 311: 1283–1287. https://doi.org/10.1126/science.1123061.

Clarridge JE (2004) Impact of 16S rRNA gene sequence analysis for identification of bacteria on clinical microbiology and infectious diseases. Clin Microbiol Rev 17: 840–862. https://doi.org/10.1128/CMR.17.4.840-862.2004.

Deurenberg RH, Bathoorn E, Chlebowicz MA, Couto N, Ferdous M, García-Cobos S, et al (2017) Application of next generation sequencing in clinical microbiology and infection prevention. J Biotechnol 243: 16–24. https://doi.org/10.1016/j.jbiotec.2016.12.022.

Devanga-Ragupathi NK, Muthuirulandi SDP, Inbanathan FY, Veeraraghavan B (2018) Accurate differentiation of Escherichia coli and Shigella serogroups: challenges and strategies. New Microbes New Infect 2: 58–62. https://doi.org/10.1016/j.nmni.2017.09.003.

Devulder G, de Montclos MP, Flandrois JP (2005) A multigene approach to phylogenetic analysis using the genus Mycobacterium as a model. Int J Syst Evol Microbiol 55: 293–302. https://doi.org/10.1099/ijs.0.63222-0.

Ibal JC, Pham HQ, Park CE, Shin JH (2019) Information about variations in multiple copies of bacterial 16S rRNA genes may aid in species identification. PLoS One 14: e0212090. https://doi.org/10.1371/journal.pone.0212090.

Janda JM, Abbott SL (2007) 16S rRNA gene sequencing for bacterial identification in the diagnostic laboratory: Pluses, perils, and pitfalls. J Clin Microbiol 45: 2761–2764. https://doi.org/10.1128/JCM.01228-07.

Johnson JS, Spakowicz DJ, Hong BY, Petersen LM, Demkowicz P, Chen L, et al (2019) Evaluation of 16S rRNA gene sequencing for species and strain-level microbiome analysis. Nat Commun 10: 1–11. https://doi.org/10.1038/s41467-019-13036-1.

Johnston PR, Quijada L, Smith CA, Baral HO, Hosoya T, Baschien C, et al (2019) A multigene phylogeny toward a new phylogenetic classification of Leotiomycetes. IMA Fungus 10, 1. https://doi.org/10.1186/s43008-019-0002-x.

Kerkhof LJ, Dillon KP, Haggblom MM, McGuinness LR (2017) Profiling bacterial communities by MinION sequencing of ribosomal operons. Microbiome 5: 116. https://doi.org/10.1186/s40168-017-0336-9.

Kumar S, Stecher G, Li M, Knyaz C, Tamura K (2018) MEGA X: Molecular evolutionary genetics analysis across computing platforms. Mol Biol Evol 8: 1829. https://doi.org/10.1093/molbev/msy096.

Lal D, Verma M, Lal R (2011) Exploring internal features of 16S rRNA gene for identification of clinically relevant species of the genus Streptococcus. Ann Clin Microbiol Antimicrob 10: 28. https://doi.org/10.1186/1476-0711-10-28.

Leray M, Knowlton N, Ho SL, Nguyen BN, Machida RJ (2019) GenBank is a reliable resource for 21^st^ century biodiversity research. Proc Natl Acad Sci USA 116: 22651–22656. https://doi.org/10.1073/pnas.1911714116.

Liu Y, Lai Q, Shao Z (2018) Genome analysis-based reclassification of Bacillus weihenstephanensis as a later heterotypic synonym of Bacillus mycoides. Int J Syst Evol Microbiol 68: 106–112. https://doi.org/10.1099/ijsem.0.002466.

Martínez-Romero E, Rodríguez-Medina N, Beltrán-Rojel M, Silva-Sánchez J, Barrios-Camacho H, Pérez-Rueda E, et al (2018) Genome misclassification of Klebsiella variicola and Klebsiella quasipneumoniae isolated from plants, animals and humans. Salud Publica Mex 60: 56–62. https://doi.org/10.21149/8149.

Mateo-Estrada V, Grana-Miraglia L, Lopez-Leal G, Castillo-Ramirez S (2019) Phylogenomics reveals clear cases of misclassification and genus-wide phylogenetic markers for Acinetobacter. Genome Biol Evol 11: 2531–2541. https://doi.org/10.1093/gbe/evz178.

Parks DH, Waite DW, Skarshewski A, Chuvochina M, Rinke C, Hugenholtz P, et al (2018) A standardized bacterial taxonomy based on genome phylogeny substantially revises the tree of life. Nat Biotechnol 36: 996–1004. https://doi.org/10.1038/nbt.4229.

Paul B, Dixi G, Murali TS, Satyamoorthy K (2019) Genome-based taxonomic classification. Genome 62: 45–52. https://doi.org/10.1139/gen-2018-0072.

Peker N, Garcia-Croes S, Dijkhuizen B, Wiersma HH, Van Zanten E, Wisselink G, et al (2019) A comparison of three different bioinformatics analyses of the 16S-23S rRNA encoding region for bacterial identification. Front Microbiol 10: 620. https://doi.org/10.3389/fmicb.2019.00620.

Quast C, Pruesse E, Yilmaz P, Gerken J, Schweer T, Yarza P, et al (2013) The SILVA ribosomal RNA gene database project: improved data processing and web-based tools. Nucleic Acids Res 41: D590–D596. https://doi.org/10.1093/nar/gks1219.

Reller LB, Weinstein MP, Petti CA (2007) Detection and identification of microorganisms by gene amplification and sequencing. Clin Infect Dis 44: 1108–1114. https://doi.org/10.1086/512818.

Sabat AJ, van Zanten E, Akkerboom V, Wisselink G, van Slochteren K, de Boer RF, et al (2017) Targeted next-generation sequencing of the 16S-23S rRNA region for culture-independent bacterial identification increased discrimination of closely related species. Sci Rep 7: 1–12. https://doi.org/10.1038/s41598-017-03458-6.

Schloss PD (2020) Reintroducing mothur: 10 Years Later. Appl Environ Microbiol 86: e02343–19. https://doi.org/10.1128/AEM.02343-19.

Sievers F, Wilm A, Dineen D, Gibson TJ, Karplus K, Li W, et al (2011) Fast, scalable generation of high-quality protein multiple sequence alignments using Clustal Omega. Mol Syst Biol 7: 539. https://doi.org/10.1038/msb.2011.75.

Srinivasan R, Karaoz U, Volegova M, MacKichan J, Kato-Maeda M, Miller S, et al (2015) Use of 16S rRNA gene for identification of a broad range of clinically relevant bacterial pathogens. PLoS One 10: e0117617. https://doi.org/10.1371/journal.pone.0117617.

Stackebrandt E, Mondotte JA, Fazio LL, Jetten M (2021) Authors need to be prudent when assigning names to microbial isolates. Arch Microbiol 203: 5845–5848. https://doi.org/10.1007/s00203-021-02599-7

Starke R, Pylro VS, Morais DK (2021) 16S rRNA gene copy number normalization does not provide more reliable conclusions in metataxonomic surveys. Microb Eco 81: 535–539. https://doi.org/10.1007/s00248-020-01586-7.

Steven B, Hesse C, Soghigian J, Gallegos-Graves V, Dunbar J (2017) Simulated rRNA/DNA ratios show potential to misclassify active populations as dormant. Appl Environ Microbiol 83: e00696–17. https://doi.org/10.1128/AEM.00696-17.

Thiergart T, Landan G, Martin WF (2014) Concatenated alignments and the case of the disappearing tree. BMC Evol Biol 14: 1–12. https://doi.org/10.1186/s12862-014-0266-0.

Varghese NJ, Mukherjee S, Ivanova N, Konstantinidis KT, Mavrommatis K, Kyrpides NC, et al (2015) Microbial species delineation using whole genome sequences. Nucleic Acids Res 43: 6761–6771. https://doi.org/10.1093/nar/gkv657.

Vetrovsky T, Baldrian P (2013) The variability of the 16S rRNA gene in bacterial genomes and its consequences for bacterial community analyses. PLoS One 8: e57923. https://doi.org/10.1371/journal.pone.0057923.

Weisburg WG, Barns SM, Pelletier DA, Lane DJ (1991) 16S ribosomal DNA amplification for phylogenetic study. J Bacteriol 173: 697–703. https://doi.org/10.1128/jb.173.2.697-703.1991.

Winand R, Bogaerts B, Hoffman S, Lefevre L, Delvoye M, van Braekel J, et al (2020) Targeting the 16S rRNA gene for bacterial identification in complex mixed samples: Comparative evaluation of second (Illumina) and third (Oxford Nanopore technologies) generation sequencing technologies. Int J Mol Sci 21: 298. https://doi.org/10.3390/ijms21010298.

Yoon SH, Ha SM, Kwon S, Lim J, Kim Y, Seo H, et al (2017) Introducing EzBioCloud: a taxonomically united database of 16S rRNA gene sequences and whole-genome assemblies. Int J Syst Evol Microbiol 67: 1613–1617. https://doi.org/10.1099/ijsem.0.001755.

